# Bet-hedging strategies in expanding populations

**DOI:** 10.1101/429506

**Authors:** Martín Paula Villa, Miguel A. Muñoz, Simone Pigolotti

## Abstract

In ecology, species can mitigate their extinction risks in uncertain environments by diversifying individual phenotypes. This observation is quantified by the theory of bet-hedging, which provides a reason for the degree of phenotypic diversity observed even in clonal populations. The theory of bet-hedging in well-mixed populations is rather well developed. However, many species underwent range expansions during their evolutionary history, and the importance of phenotypic diversity in such scenarios still needs to be understood. In this paper, we develop a theory of bet-hedging for populations colonizing new, unknown environments that fluctuate either in space or time. In this case, we find that bet-hedging is a more favorable strategy than in well-mixed populations. For slow rates of variation, temporal and spatial fluctuations lead to different outcomes. In spatially fluctuating environments, bet-hedging is favored compared to temporally fluctuating environments. In the limit of frequent environmental variation, no opportunity for bet-hedging exists, regardless of the nature of the environmental fluctuations. For the same model, bet-hedging is never an advantageous strategy in the well-mixed case, supporting the view that range expansions strongly promote diversification. These conclusions are robust against stochasticity induced by finite population sizes. Our findings shed light on the importance of phenotypic heterogeneity in range expansions, paving the way to novel approaches to understand how biodiversity emerges and is maintained.

**Author summary:** Ecological populations are often exposed to unpredictable and variable environmental conditions. A number of strategies have evolved to cope with such uncertainty. One of them is stochastic phenotypic switching, by which some individuals in the community are enabled to tackle adverse conditions, even at the price of reducing overall growth in the short term. In this paper, we study the effectiveness of these “bet-hedging” strategies for a population in the process of colonizing new territory. We show that bet-hedging is more advantageous when the environment varies spatially rather than temporally, and infrequently rather than frequently.

## Introduction

The dynamics and evolutionary history of many biological species, from bacteria to humans, are characterized by invasions and expansions into new territory. The effectiveness of such expansions is crucial in determining the ecological range and therefore the success of a species. A large body of observational [1, 2] and experimental [3–6] literature indicates that evolution and selection of species undergoing range expansions can be dramatically different from that of other species resident in a fixed habitat. Theoretical studies of range expansions based on the Fisher-Kolmogorov equation [7, 8] or variants [9, 10] also support this idea. Adaptive dispersal strategies [2] and small population sizes at the edges of expanding fronts [11, 12] are among the main reasons for such differences.

Range expansions often occur in non-homogeneous and fluctuating environments. Under such conditions, it is possible to mathematically predict the expansion velocity of a community of phenotypically identical individuals [13–18]. However, diversity among individuals is expected to play an important positive role when populations expand in fluctuating environments. For instance, diverse behavioral strategies help animal populations to overcome different invasion stages and conditions [19–22]. Analyses of phenotypic diversity in motile cells suggest that it also may lead to a selective advantage at a population level [23–25]. Although several studies have tackled the problem of how individual variability affects population expansion [6, 9, 10, 26, 27, 27, 28, 28–30], systematic and predictive theory is still lacking [22].

Phenotypic diversification is often interpreted as a bet-hedging strategy, spreading the risk of uncertain environmental conditions across different phenotypes adapted to different environments [31–40]. Since its formalization in the context of information theory and portfolio diversification [41, 42], a large number of works have explored the applicability of bet-hedging in evolutionary game theory [43–46] and ecology [47–51]. Few studies have explored the benefits of bet-hedging in spatially structured ecosystems [52, 53].

In this paper, we study how bet-hedging strategies can aid populations in invading new territories characterized by fluctuating environments. In particular, we analyze the effect of spatial expansion, different types of environmental heterogeneity, and demographic stochasticity on development of bet-hedging strategies for a population front evolving according to a Fisher wave.

By employing mathematical as well as extensive computational analyses, we find that the advantage of bet-hedging in range expansions depends on the rate of environmental variation. In particular, bet-hedging is more convenient for infrequently varying environments, whereas its advantages vanish for frequent environmental variation. For the same model, bet-hedging is never an advantageous strategy in the well-mixed case, supporting the view that range expansions strongly promote diversification. We further find that spatial environmental variations provide more opportunities for bet-hedging than temporal fluctuations. Finally, we show that our conclusions still hold when considering stochastic effects on the front propagation induced by a finite population size.

The paper is organized as follows. In Section we introduce a general population model and an example with two available phenotypes and two environmental states. Section presents an extensive study of this example. In, we demonstrate that the main conclusions obtained for the example also hold for the general model. Section is devoted to conclusions and perspectives.

## Results

### Model

We consider a population consisting of individuals that can assume *N* alternative phenotypes. The population as a whole adopts a phenotypic strategy, that is identified by the fractions *α_i_*, *i* = 1… *N* of the population assuming each phenotype *i* with ∑_*i*_*α_i_* = 1 and 0 ≤ *α_i_* ≤ 1 ∀*i* (Fig. 1A). As customary in game theory, we say that a strategy is a “pure strategy” if *α_i_* = *δ_ik_* for some phenotype *k*, and a “mixed strategy” otherwise. We assume that the *α_i_*’s remain constant in time within the population.

**Fig 1.**
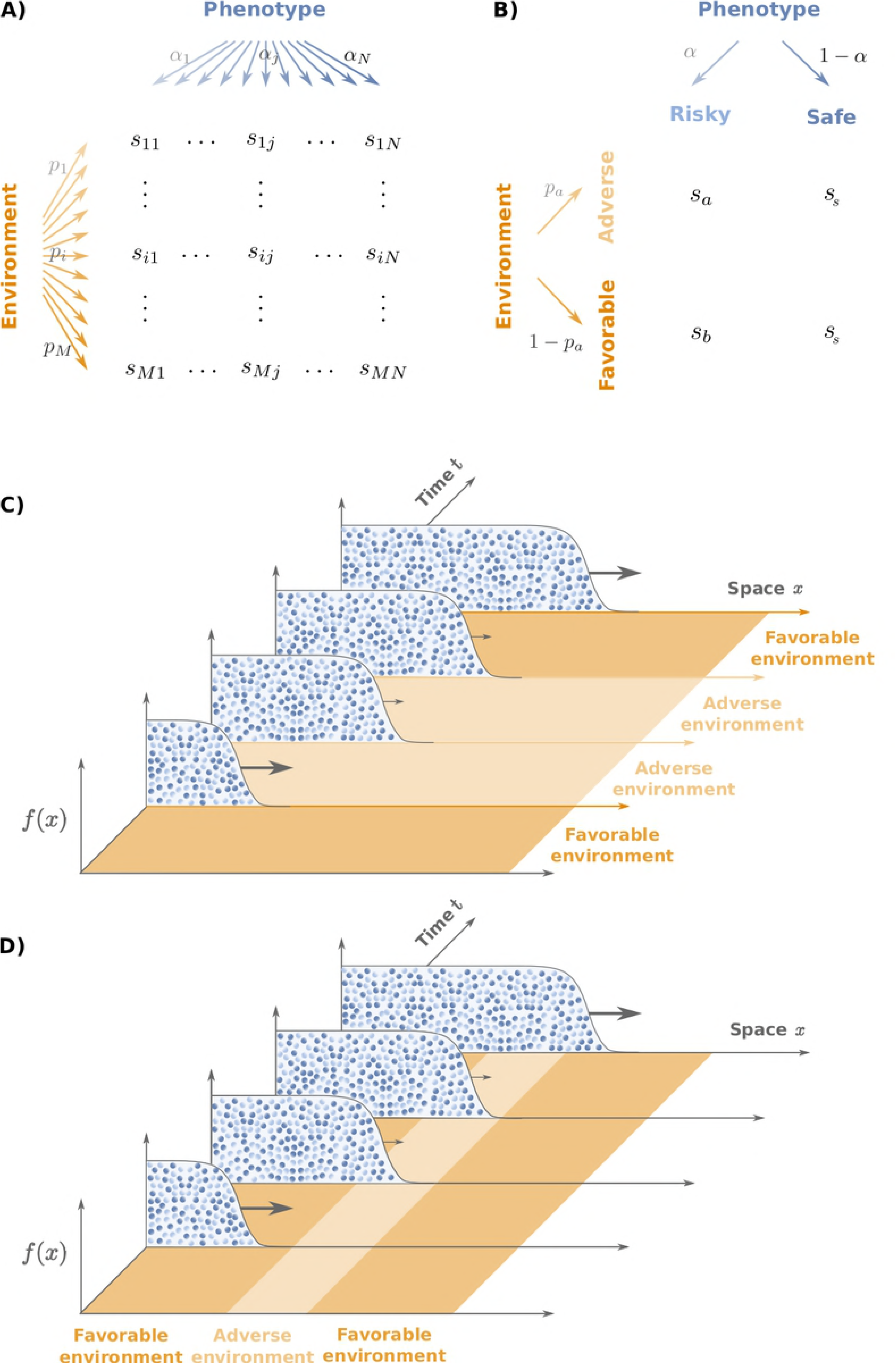
Population model. A) General model: individuals can adopt N different phenotypes with probabilities *α_j_*(*j* = 1, …, *N*) and experience *M* different environmental conditions with probabilities *p_i_* (*i* = 1, …,*M*). The fitness of an individual with phenotype *j* in an environment *i* is given by *S_ij_*. B) Two-phenotypes model: Individuals can adopt either a “risky” or a “safe” phenotype with probabilities *α*, and 1 —*α* respectively. The safe phenotype is characterized by an environment-independent growth rate *s_s_*. The growth rate of the risky phenotype is *s_a_* or *s_b_*, depending on whether the current environment is “adverse” (a) or “favorable” (b). C) and D) Sketch of range expansion in a population having 0 ≤ *α* ≤ 1 for temporally varying C) and spatially varying D) environments, respectively.

The environment can be found in one of *M* different states, which can randomly alternate either in time or in space. We call *p_i_* the probability of encountering environment *i*. We further define the growth rate *s_ij_* ≥ 0 of phenotype *j* in environment *i* (Fig. 1A). When the population size is sufficiently large, so that demographic stochasticity can be neglected, the population-averaged growth rate given the state *i* = *i(x,t)* of the environment at position *x* and time *t* is 
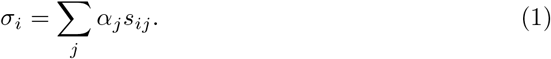

Since Eq. (1) is linear in the *α_j_*’s, the population-averaged growth rate in a given environment is always maximized by the pure strategy with the highest growth rate. However, in the presence of uncertainty about the environment, the population might choose other strategies. One possibility is to select a different pure strategy, less risky than the optimal one. This case is often termed “conservative bet-hedging” in the ecological literature [40]. Another option is to adopt a mixed strategy, with different phenotypes more adapted to different environments. This case is termed “diversifying bet-hedging” in the literature [40, 54]. Since our interest is in diversification, the term “bet-hedging” will refer herein to diversifying bet-hedging.

Before presenting our results in full generality, we will illustrate it in a simple, yet ecologically relevant instance of the model with only two phenotypes: “safe” and “risky” and two environmental states: “adverse” (a) and “favorable” (b). The safe phenotype is characterized by a growth rate *s_s_* both in the adverse and favorable environments. The growth rate of the risky phenotype is *s_a_* in environment (a) and *s_b_* in environment (b) (Fig. 1B) [55]. The two environments occur with the same probability, *p_a_* = *p_b_* = 1/2. A fraction of individuals *α* adopts the risky phenotype and the complementary fraction (1 − *α*) adopts the safe phenotype (Fig. 1B). For this model, the population-averaged growth rate reads 
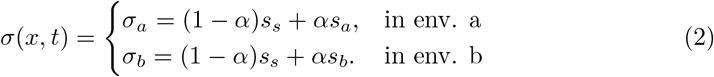

Note that, with a slight abuse of notation, we use equivalently σ*_i_* or σ(*x,t*) to denote the population-averaged growth rate in the environment *i*(*x,t*). For pure strategies, *α* = 0 or *α* = 1, the population-averaged growth rate *σ* reduces to the growth rate of the safe or risky phenotype, respectively.

### Two-phenotype model

We seek to understand those conditions under which bet-hedging is advantageous for the population. To this end, we shall compare three situations: i) well-mixed populations, ii) range expansions in environments that fluctuate temporally, but that are homogeneous in space (Fig. 1C), and iii) range expansions in spatially fluctuating environments that are homogeneous in time (Fig. 1D).

### Well-mixed case

We start by analyzing the well-mixed case, where the spatial coordinates of individuals can be ignored. The total population density *f* (*t*) evolves according to the equation 
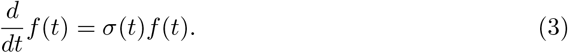

In writing Eq. (3), we used the assumption that the fraction *α* of the population adopting the risky phenotype remains constant in time (see [56, 57] for cases in which this assumption is relaxed). Equation (3) can be readily integrated, obtaining 
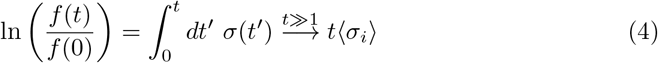
where 〈σ*_i_*〉 = ∑_*i*_*p_i_*σ_*i*_ denotes an average over the environmental states. For Eq. (4) to hold, we do not need to make strong assumptions about the statistics of the environmental states, other than it should be stationary, ergodic, and with a finite correlation time.

The optimal strategy *α*^*^ is obtained by maximizing the right-hand side of Eq. (4) respect to the strategy *α*. Since 〈 σ*_i_*〉 is a linear function of *α*, its maximum is always reached at the extremes of the interval (*α* ∊ [0,1]). In particular, defining the normalized growth rates 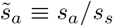 and 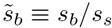, we find that the optimal strategy is *α\** = 1 when 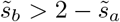 and *α*^*^ = 0 otherwise. This means that no bet-hedging strategy is possible in this model in the well-mixed case [55].

This simple result illustrates an aspect of bet-hedging that is sometimes under-appreciated. In well-mixed systems, bet-hedging requires at least one of the following ingredients: a) discrete generations, as in the seminal work of Kelly [41], b) finite switching rates among strategies [32, 57], or c) a delta-correlated environment [52]. Any of these ingredients can lead to nonlinearities in the average exponential growth rate, therefore opening the way for a non-trivial optimal strategy. A detailed analysis of these facts is beyond the scope of the present work and will be presented elsewhere.

Note that, in this model, the frequency of environmental change does not play a role, as far as it is finite [52]. The physical reason can be understood from the right-hand side of Eq. (4): the optimal strategy depends on the frequency of different environmental states but not on the switching rates. This feature is also shared by other well-mixed models that do allow for optimal bet-hedging strategies, such as the classic model by Kelly [41]. We shall see in the following that, on the contrary, the rate of environmental change plays an important role for expanding populations.

### Range expansion in fluctuating environments

We now consider a population expanding into an unoccupied, one-dimensional space under the influence of a stochastically changing environment. Its population dynamics are described by the Fisher equation [7, 58]:
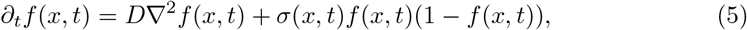
where *f* (*x, t*)is the population density at spatial coordinate *x* and time t, and *D* is the diffusion constant, which characterizes the motility of individuals. For a constant growth rate σ, the stationary solution of Eq. (5) is characterized by a front advancing in space with velocity 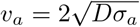. Instead, we consider a fluctuating case in which the growth rate σ(*x,t*) depends on the population strategy *α* and on environmental conditions according to Eq. (2). In such case, one can define an asymptotic mean velocity of the front as 
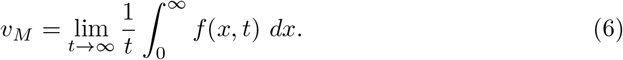

In what follows, we take *v_M_* as a proxy of the long-term population fitness and maximize it with respect to *α* to determine the optimal strategy.

### Range expansion in temporally varying environments

We first consider the case in which environmental conditions change randomly with time, but are homogeneous across space, σ(*x,t*) =σ(*t*)(see Fig.1C). Switching rates between adverse and favorable environments are *k_a_*_→_*_b_* = *k_b_*_→*a*_ = *k*. We first estimate the asymptotic mean velocity defined in Eq. (6) in the limiting cases of *k* → 0 and *k*→∞.

When the environment changes very infrequently, *k*→0, the population front has the time to relax to the asymptotic shape characterized by its corresponding Fisher velocity, 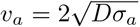 or 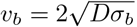 depending on the environment [7, 59]. Thus, the asymptotic mean velocity can be estimated as *v_M_* = (*v_a_* + *v_b_*)/2. Maximizing *v_M_* with respect to *α*, we find that in this case, a bet-hedging optimal strategy exists under the conditions (Fig. 2A): 
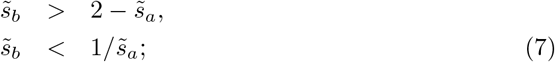
observe that the first condition coincide with the one in the well-mixed scenario. In the opposite limiting case of a rapidly fluctuating environment, *k*→ ∞, the population effectively experiences the average of the two growth rates, so that the velocity can be estimated as 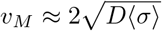, where 〈(…)〉 denotes an average over the environmental states. In this case, the optimal strategy *α*^*^ is achieved by maximizing the average growth rate 〈σ〉 with respect to *α*. Since 〈σ〉 is linear in *α*, the maximum always lies at the extremes of the interval [0,1]. In particular, we find *α*^*^ = 1 when 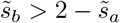 and *α*^*^ = 0 otherwise, as in the well-mixed case. This implies that no bet-hedging regime exists in this limit (Fig. 2B).

**Fig 2.**
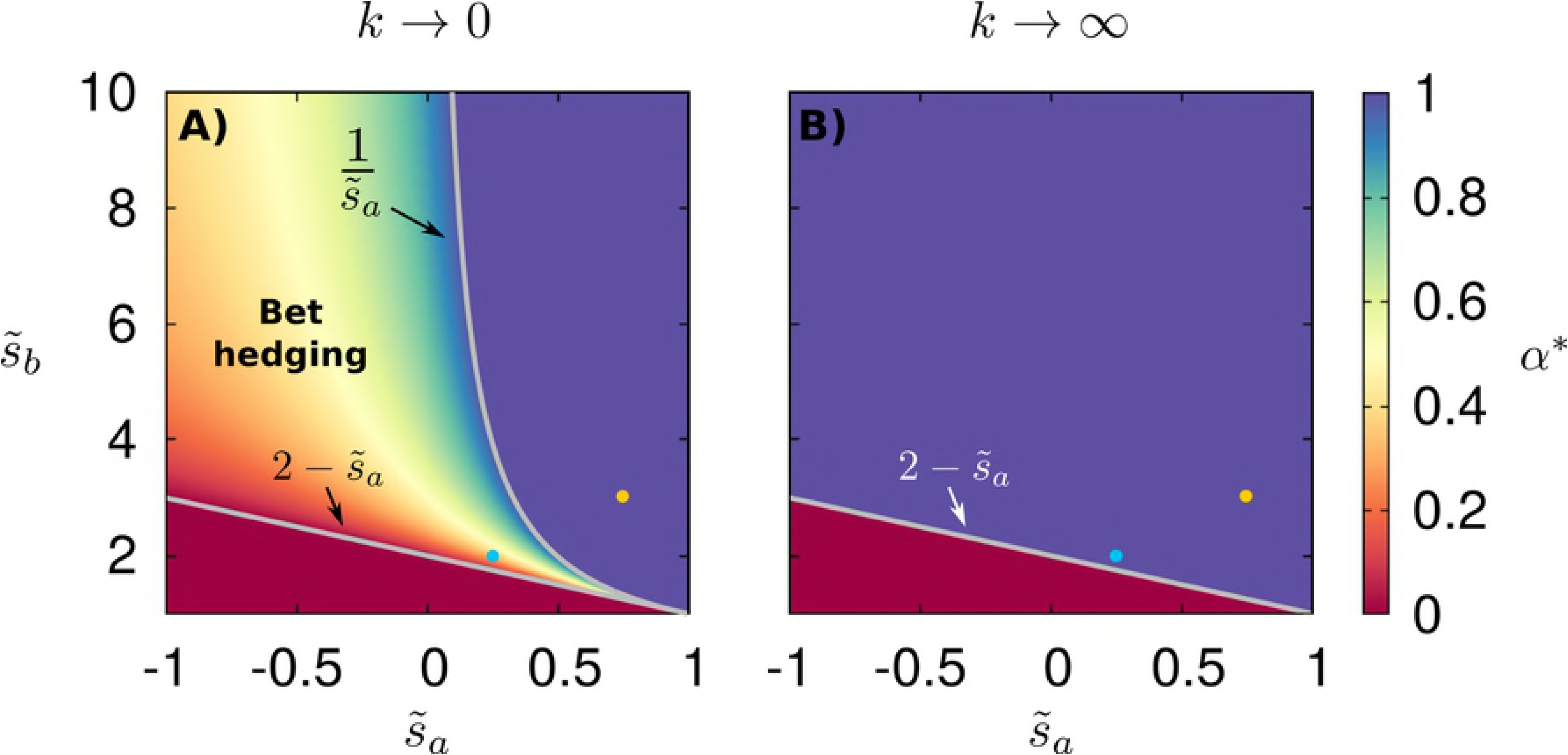
Bet-hedging region in temporally varying environments. Optimal strategy *α*^*^ as a function of growth rates 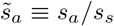 and 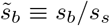 for range expansions in temporally varying environments under the limits of environmental change rate (A) *k* → 0, see Eq.(7), and (B) *k* → ∞. In all panels, lines delimit the bet-hedging region 0 ≤ *α*^*^ ≤1. Two dots in the panels mark parameter values chosen for the analysis of Figs. 3,4,5.

**Fig 3.**
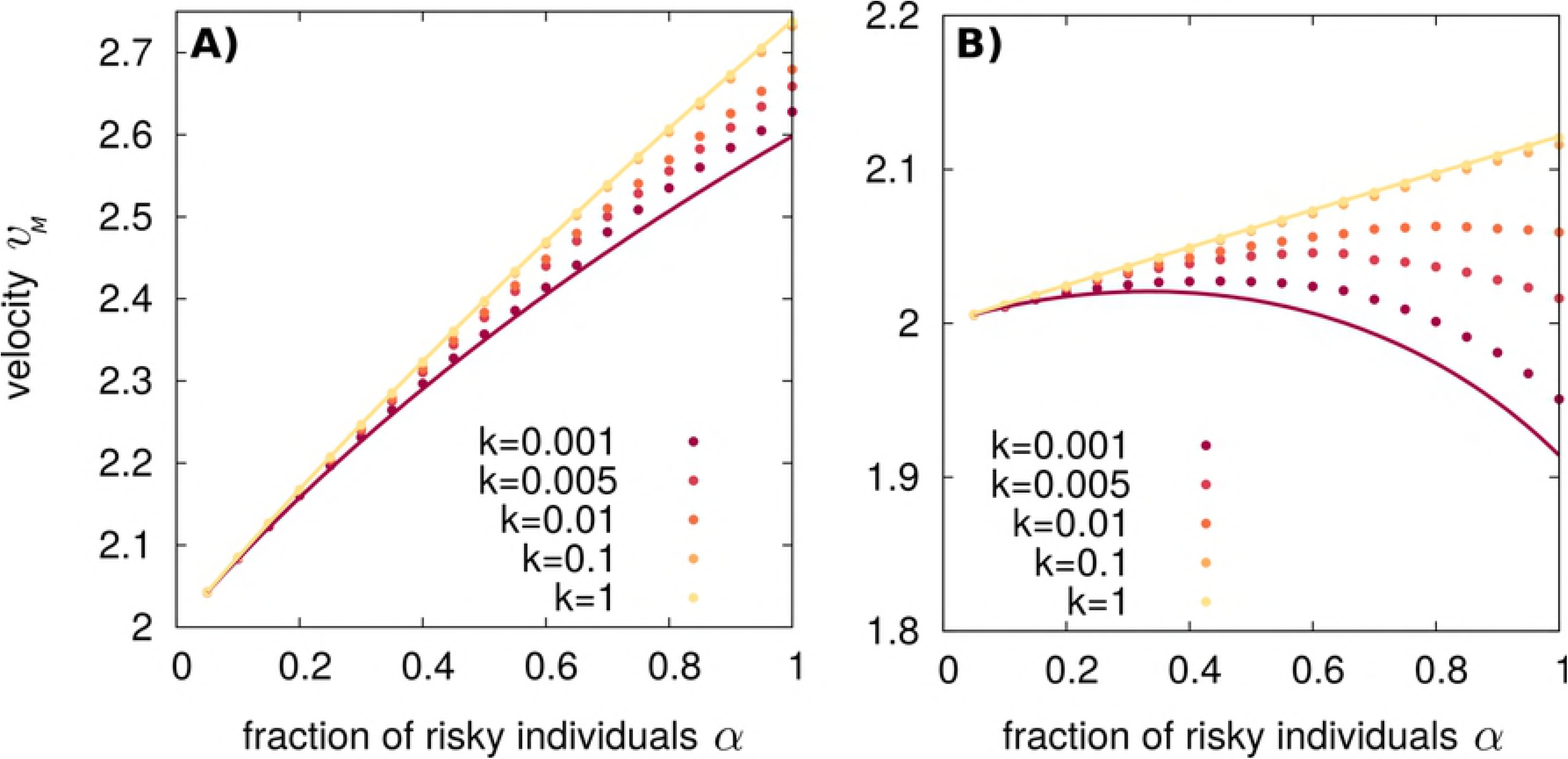
The asymptotic mean velocity increases with *k* in temporally varying environments. (A) Velocities obtained by numerical integration of Eq. 5 for *s_a_* = 0.75, *s_s_* = 1, *s_b_* = 3 (yellow dot of Fig. 2) for different switching rates *k* shown in the figure legend. (B) The same for *s_a_* = 0.25, *s_s_* = 1, *s_b_* = 2 (blue dot of Fig. 2). In (A), the optimal strategy is *α* = 1 for all *k* values. In (B), bet-hedging optimal strategies appear depending on the value of *k*. The continuous red and yellow lines (both in A and B) illustrate analytical predictions under the two limits *v_M_*(*k*→0)= (*v_a_*(*α*)+*v_b_*(*α*))/2 and 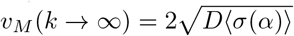, respectively.

**Fig 4.**
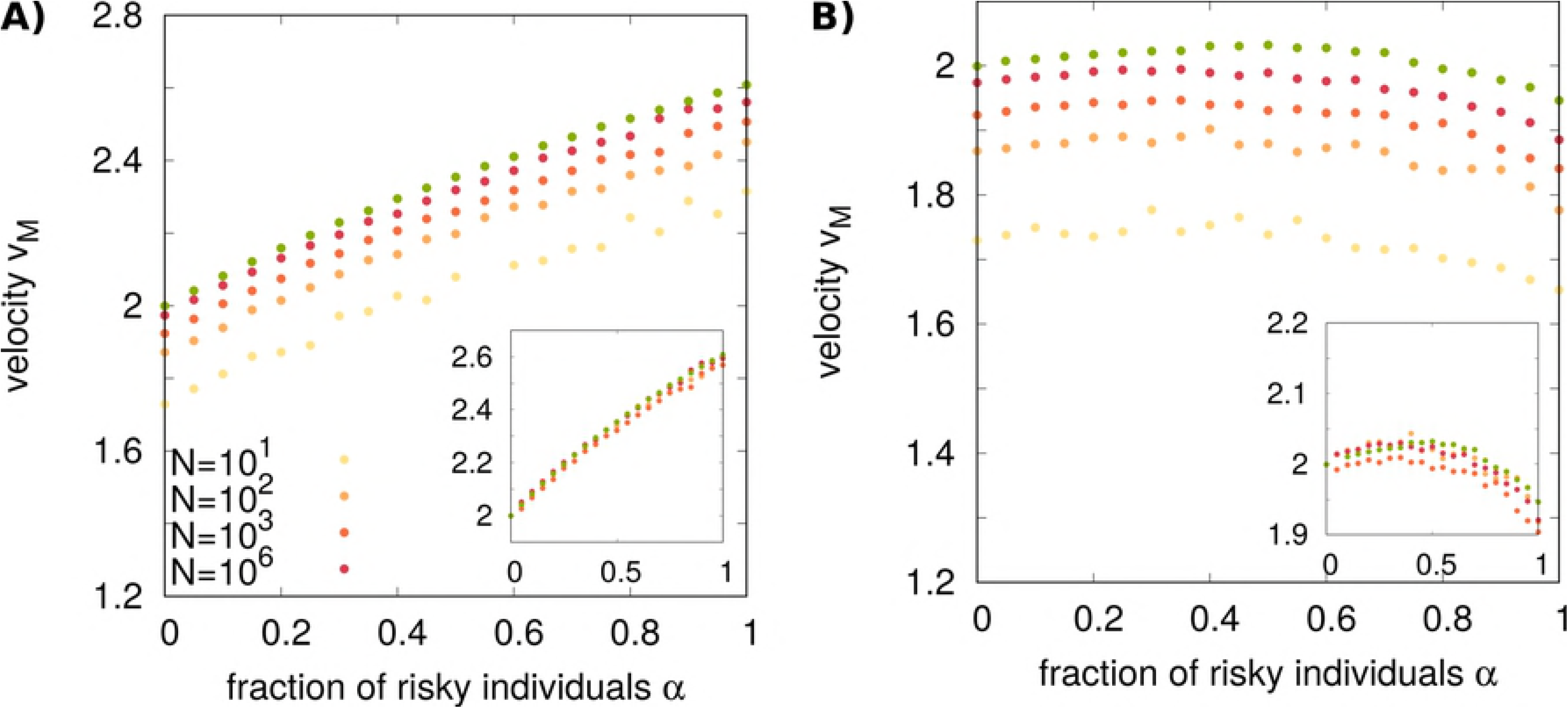
The bet-hedging region is expanded for range expansions in spatially varying environments compared to temporally varying environments. A)Optimal strategy *α*^*^ as a function of the parameters for spatially varying environments in the limit *k_s_* → 0, Eq. (9). White lines mark the limits of the bet-hedging region. The limit for which the strategy *α* = 1 is optimal in temporally fluctuating environments for *k* → 0 is also shown (gray line) for comparison. B) The velocity obtained by numerical integration of Eq. (5) for *s_a_* = 0.25, *s_s_* = 1, *s_b_* = 2 (corresponding to the blue dot of panel A) and different values of *k_S_* shown in the figure legend. Light and dark gray lines correspond to the analytical limits for temporally varying environments,*v_M_*(*k*→0) = (*v_a_*(*α*)+*v_b_*(*α*))/2, and 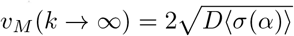, respectively. The red curve is the analytical solution for a spatially fluctuating environment with *k_S_* → 0, see Eq. (9). Note that in this case, the asymptotic mean velocity does not increase monotonically with *k_S_* but is maximal at *k_s_* ≈ 0.1.

**Fig 5.**
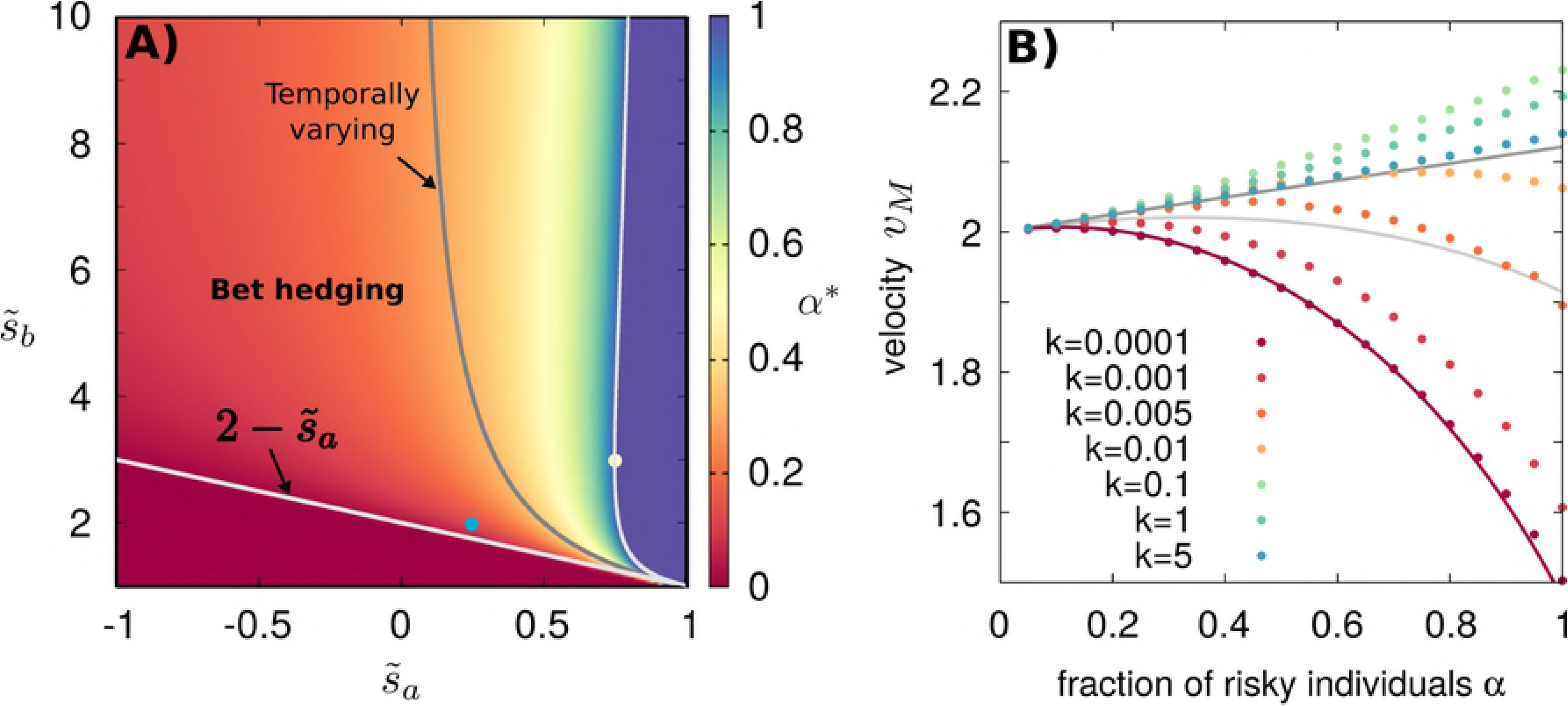
The optimal strategy is robust with respect to noise induced by finite population size in temporally varying environments. (A) Asymptotic mean velocities obtained by numerical integration of the stochastic Fisher equation (10) for 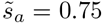, *s_s_* = 0.01, 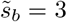 (yellow dot of Fig. 2) and different population sizes. (B) The same for 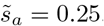, *s_s_* = 1, 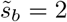 (blue dot of Fig. 2). In both panels, the temporal switching rate of the environment is k = 0.001. Green dots corresponds to the results of Figs. 3A,B for *k* = 0.001. Insets show a collapse of the curves according to Eq. (11), with a fitted value of *C* = 3.

To explore the intermediate regimes of finite *k*, it is necessary to resort to numerical simulations of Eq. (5). For a set of parameters such that the optimal strategy is *α*_*_ = 1 for *k* → 0, the optimal strategy remains *α*_*_ = 1 for all values of *k*, see Fig. 3A. Instead, in a case where the optimal solution is in the bet-hedging region for *k* → 0, the optimal strategy *α*_*_ increases with the switching rate, so that for large *k* the optimal strategy is outside the bet-hedging region, *α*_*_ = 1. These results support our analytical estimates of limiting values and suggest that the asymptotic mean velocity is a monotonically increasing function of the switching rate *k* in this case.

### Range expansion in spatially varying environments

We now consider the case in which environmental conditions are constant in time, but depend on the spatial coordinate *x*. The dynamics are described by the Fisher equation (5) with two types of environment randomly alternating in space, σ(*x,t*) = σ(*x*). We call *k_S_* the spatial rate of environmental switch, so that the probability of encountering an environmental shift within an infinitesimal spatial interval *dx* is equal to *k_S_dx*. The switching rates from environment *a* to *b* and vice-versa are both equal to *k_S_*. As above, we first analyze the two limits *k_S_* → 0 and *k_S_* → ∞.

In the limit *k_S_* → 0, the population front traverses large regions of space characterized by a constant environment, either *a* or *b*, thus being able to reach the corresponding Fisher velocity, *v_a_* or *v_b_*, respectively. The mean traversed lengths Δ*x_a_* and Δ*x_b_* are equal for the two environments. On the other hand, the mean times spent in each of them, *t_a_* and *t_b_*, are different, and satisfy the relation 
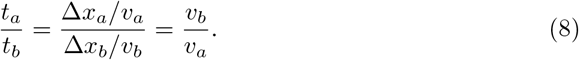

Therefore, in this case, the asymptotic mean velocity is given by the harmonic mean of the velocities in the two environments 
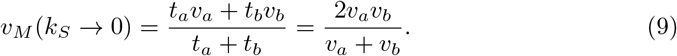

At the opposite limit of large *k_S_*, the environment is characterized by frequent spatial variations. In this case, the population front occupies multiple *a* and *b* sectors with an effective growth rate 〈σ〉. As in the time-varying case, the asymptotic mean velocity in this limit is, 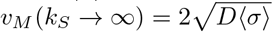, see also [14, 15].

Here, for *k_S_* → 0 the bet-hedging region is broader with respect to the temporally fluctuating environment for *k* → 0, see Fig. 4A. For *k_S_* → ∞, the optimal strategy is the same as in Fig. 2C and there is no bet-hedging regime.

We numerically solved Eq. (5) for intermediate values of *k_S_* and obtained the mean asymptotic velocities as a function of *α*, see Fig. 4 B. Results support theoretical predictions in the limiting cases *k_S_* → 0 and *k_S_* → ∞. In this case, we observe a non-monotonic behavior of *v_M_* as a function of *k_S_*, so that the maximum mean velocity is attained at an intermediate switching rate. An analytical explanation of this non-trivial effect goes beyond the scope of this work.

### Effect of finite population size

The deterministic Fisher equation (5) is rigorously valid only in the limit of infinite local population sizes. We now explore the robustness of our results when considering stochasticity induced by the finite size of populations, i.e. “demographic noise”. We focus on the case of a front propagating in a temporally varying environment. To study finite population size, we solve numerically a stochastic counterpart of the Fisher equation 
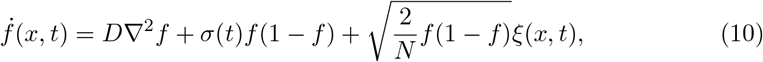
see e.g. [60]. In Eq. (10), ξ(*x,t*) is Gaussian white noise with 〈ξ(*x,t*)〉 = 0,〈ξ(*x,t*)ξ(*x′*, *t′*)〉 = δ(*x*−*x′*)δ(*t*−*t′*). The parameter *N* represents the number of individuals per unit length corresponding to *f*(*x, t*) = 1. For large population sizes, *N* ≫ 1, Eq. (10) reduces to Eq. (5). Numerical integration of Eq. (10) requires some care due to the fact that both noise and the deterministic terms go to zero as the absorbing states *f*(*x,t*) = 0 and *f*(*x,t*) = 1 are approached [61, 62]. A detailed description of our integration scheme is presented in the Supporting information.

For a Fisher wave propagating in a homogeneous environment, demographic noise leads to a reduced front velocity *v* with respect to the deterministic case [58, 62–64]
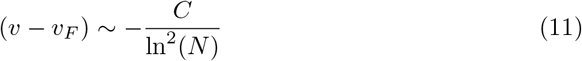
where *C* is a constant, *N* is the maximum population size per unit length, and *s_s_* = 1, 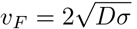 is the Fisher velocity in the absence of demographic noise. Asymptotic mean velocities for stochastic waves in temporally varying environments are shown in Fig. 5. Also in this case, small populations, subject to relatively strong demographic noise, propagate more slowly than large populations. In particular, curves at different values of *N* can be approximately rescaled using Eq. (11), assuming that *C* does not depend on *α* (insets of Fig. 5). These results imply that the optimal strategy *α*^*^ is robust with respect to demographic noise, at least for moderately to relatively large values of *N*. The same scaling holds for spatially varying environments, but with mild deviations that seem to expand the bet-hedging region even further, compared with the infinite population size limit (see Supporting information).

### General bet-hedging model

In this section, we demonstrate that our main conclusions hold also for the general case with *N* phenotypes and *M* environmental states (see Section). In particular, for a temporally fluctuating environment in the limit of very slow switching rates, the bet-hedging regime occupies a reduced region of parameter space compared to temporally constant environments fluctuating slowly in space. Also in this case, we find that for frequent environmental change, the propagation velocity tends to *s_s_* = 1, 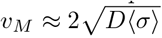, regardless of whether the environmental fluctuations depend on time or space. Therefore, the optimal strategy maximizes the linear function of the *α_i_*S 〈σ〉 and is therefore a pure strategy as discussed after Eq. (1).

We consider a range expansion where the environment fluctuates in time and the stochastic switching rates among the *M* environmental states are small. Following the same line of thought of Section, the optimal strategy maximizes 
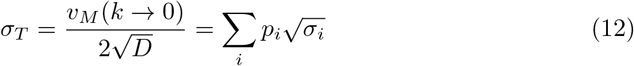
where, as usual, σ_*i*_ = ∑*_jsij_α_j_*. For spatially varying environments, the optimal strategy maximizes the harmonic mean 
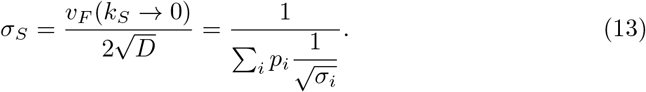

Both for Eq. (12) and Eq. (13), maximization has to be performed with the constraint ∑*_j_α_j_* = 1 and 0 ≤*α_j_*≤ 1 ∀*j*. We recall that the bet-hedging regime is the region of parameter space where the optimal solution is a mixture of all phenotypes, *α_i_*> 0 ∀*i*. Here we show that if, for a given choice of the *s_ij_*’s and *p_i_*’s, a population advancing in a temporally varying environment is in a bet-hedging regime, then the same holds for spatially varying environments. For the demonstration, we borrow a mathematical tool from evolutionary game theory [65]. We introduce the gradients 
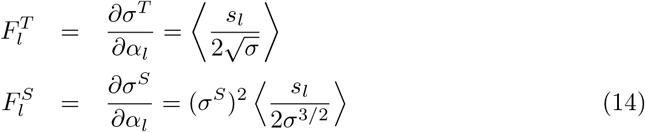
where 〈*x*〉 =∑_*i*_*p_i_x_i_* is the average over environments. We now associate replicator equations to Eq. (12) and Eq. (13): 
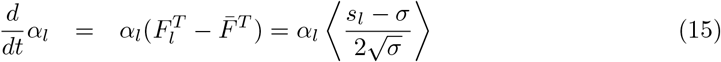

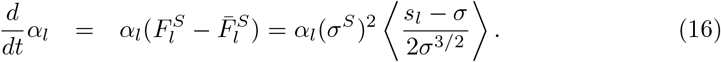

The system is in a bet-hedging regime when the replicator equations admit a stable fixed point in the interior of the unit simplex, 0 < *α_i_* < 1. Instead of computing the fixed point explicitly, we check whether each phenotype *l* has a positive growth rate for *α_l_* ≪ 1. Brouwer’s fixed point theorem ensures that, under this condition, there must be a fixed point in the interior, see [65], chapter 13. For our aims, it is therefore sufficient to prove that, for small *α_l_*, if 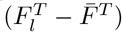 is positive, then 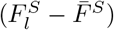 must be positive as well. Note that for *α_l_* ≪ 1, the average σ = ∑_*j*_ *s_ij_α_j_* does not depend on *α_l_*, and therefore, σ and *s_l_* are uncorrelated random variables respect to the average over the environment. Since 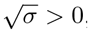, this means that the sign of 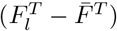 is the same than the quantity 
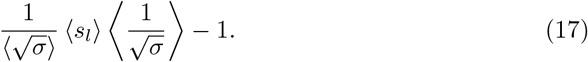

Following the same logic, the sign of 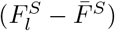 is the same than 
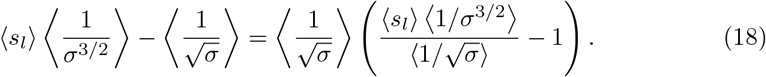

Since also 〈*s_l_*〉 > 0, we need to demonstrate that the following inequality always holds 
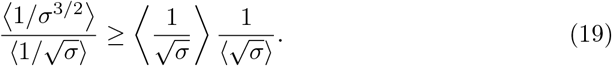

This can be proven from the chain of inequalities 
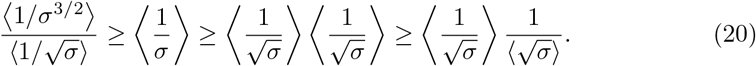

In Eq. (20), the second and third inequalities are consequences of Jensen’s inequality, since both *x*^2^ and 1/*x* are convex functions. For the first inequality in Eq. (20), since *s* > 0, we can use the result 〈*x^i^*〉 ≥ 〈*x^j^*〉 ^*i/j*^ proved for *i* > *j* in [66]. Combining this result for (*i* = 3, *j* = 2) and (*i* = 2, *j* = 1), we obtain 〈*x*^3^〉 ≥ 〈*x*^2^〉) 〉*x*〈. Taking 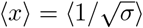 we finally prove Eq. (20). Therefore, in the limit of small switching rates of the environment, the bet-hedging region is wider in the spatially varying case than in the temporally varying case.

In the opposite limit of high rates of environmental switch, the function to be optimized is linear, and the optimal strategy is a pure strategy. In this case, the particular phenotype *l* adopted by the whole population is that maximizing ∑_*i*_*p_i_s_il_*. This conclusion holds both for temporally and spatially varying environments.

## Conclusions

Understanding the precise mechanisms of population expansions is of utmost importance, not only for understanding species diversity, but also to cope with invasive species in new habitats [19–22], bacterial infections [23–25, 67], and cell migration, such as those occurring during tissue renewal or cancer metastasis [5]. Phenotypic diversity is a convenient strategy for the success of population expansions in a broad range of contexts [19–25]. Although precise experimental measures are not easy to obtain, a recent study shows that populations with increased variability in individual risk-taking can colonize wider ranges of territories [68].

In this work, we proposed a general mathematical and computational framework to analyze such scenarios. In particular, we introduced a population model with diverse phenotypes that perform differently depending on the type of environment. We focused on the “optimal” degree of diversity leading to the fastest average population expansion in an environment fluctuating either in space or in time. We found that, contrarily to the well-mixed case, bet-hedging can be convenient in expanding populations. This result complements the study in [52] for a fixed habitat and supports the view that diversification is of broad importance for spatially-structured populations. For environments varying slowly in time, the expansion is relatively slow, and diverse communities can be optimal depending on the parameters. On the contrary, for fast environmental changes, the optimal population always adopts a unique strategy.

A remarkable outcome of our analysis is that spatial fluctuations create more opportunities for bet-hedging than temporal fluctuations, in that the region of parameter space where the optimal population is diverse, is always larger in the former case. One intuitive explanation is that in the case of spatial fluctuations, the population spends less time traversing favorable patches than adverse ones. This means that the beneficial effect of favorable patches is reduced with respect to the case of temporal fluctuations. Therefore, a pure risky strategy is less efficient in the case of spatial variability and can be more easily outcompeted by a diversified bet-hedging strategy.

The framework presented here can be extended to accommodate other scenarios. We have assumed that the fraction of individuals adopting each phenotype is fixed by the phenotypic switching rates. To understand the evolution of bet-hedging, it could be interesting to study scenarios in which the phenotypic switching rates are slower, so that phenotypes can be selected, and/or are themselves subject to evolution and selection [55, 69]. Another potentially relevant extension would be to consider two-dimensional habitats. Although the classic theory for Fisher waves [7, 8] is unaffected in higher dimensions, in the presence of spatial heterogeneity the front shape can become anisotropic, potentially affecting the results. Similarly, it would be interesting to analyze the combined effect of spatial and temporal variability. We also limited ourselves to the case where the different environments affect individual growth rates, whereas in general, one could also expect them to have an effect on motility [13, 14, 70–72], opening the way for different forms of bet-hedging. Finally, the present study was limited to pulled waves. It would be interesting to study the effect of bet-hedging on pushed waves, for example to describe population expansion in the presence of an Allee effect [73, 74].

It would be also interesting to experimentally test our results. Experiments of expanding bacterial colonies in non-homogeneous environments have already been performed and shed light, for example, on the evolution of antibiotic resistance in spatially-structured populations [75]. To perform experiments within the limits of our theory, a challenge can be to maintain the environmental variability sufficiently low to avoid exposing the population to an excessive evolutionary pressure. Similar problems appear, for example, in studies of range expansion of mutualistic bacteria [76]. An extension of the theory including both phenotypic and genetic diversity could account for these scenarios.

In summary, we have introduced a mathematical/computational model to understand conditions favoring diversification of an expanding population. Our work provides a bridge between the theory of bet-hedging and that of ecological range expansion described by reaction-diffusion equations. The results of the model highlight the relation between population diversity and fluctuations of the environment encountered during range expansion. The flexibility and generality of our framework make it a useful starting point for applications to a wide range of ecological scenarios.

## Supporting information

**S1 Appendix**. Here we provide several supplemental appendices: **Numerical integration of the Stochastic Fisher equation** where we describe in detail the methods applied for the integration of the wave equations of the two-phenotype model studied. **Effect of finite population size for spatially varying environments** where we study the effect of demographic stochasticity induced by the finite size of the population for spatially varying environments. (PDF)

## Acknowledgments

We acknowledge Steven D. Aird, R. Rubio de Casas, and Massimo Cencini for comments on a preliminary version of this manuscript. MAM is grateful to the Spanish-MINECO/AEIa for financial support (under grant ref. FIS2017–84256-P; FEDER funds).

## References

1. Ramachandran S, Deshpande O, Roseman CC, Rosenberg NA, Feldman MW, Cavalli-Sforza LL. Support from the relationship of genetic and geographic distance in human populations for a serial founder effect originating in Africa. Proceedings of the National Academy of Sciences of the United States of America. 2005;102(44):15942–15947.

2. Duckworth RA. Adaptive dispersal strategies and the dynamics of a range expansion. The American Naturalist. 2008;172(S1):S4-S17.

3. Wolfe AJ, Berg HC. Migration of bacteria in semisolid agar. Proceedings of the National Academy of Sciences. 1989;86(18):6973–6977.

4. Hallatschek O, Hersen P, Ramanathan S, Nelson DR. Genetic drift at expanding frontiers promotes gene segregation. Proceedings of the National Academy of Sciences. 2007;104(50):19926–19930.

5. Mayor R, Etienne-Manneville S. The front and rear of collective cell migration. Nature reviews Molecular cell biology. 2016;17(2):97.

6. Fu X, Kato S, Long J, Mattingly HH, He C, Vural DC, et al. Spatial self-organization resolves conflicts between individuality and collective migration. Nature communications. 2018;9(1):2177.

7. Fisher RA. The wave of advance of advantageous genes. Annals of Human Genetics. 1937;7(4):355–369.

8. Kolmogorov A, Petrovskii I, Piskunov N. A study of the diffusion equation with increase in the amount of substance, and its application to a biological problem. Selected Works of AN Kolmogorov I. 1937; p. 248–270.

9. Neubert MG, Caswell H. Demography and dispersal: calculation and sensitivity analysis of invasion speed for structured populations. Ecology. 2000;81(6):1613–1628.

10. Bartoń K, Hovestadt T, Phillips B, Travis J. Risky movement increases the rate of range expansion. Proceedings of the Royal Society of London B: Biological Sciences. 2012;279(1731):1194–1202.

11. Waters JM, Fraser CI, Hewitt GM. Founder takes all: density-dependent processes structure biodiversity. Trends in ecology & evolution. 2013;28(2):78–85.

12. Hallatschek O, Nelson DR. Gene surfing in expanding populations. Theoretical population biology. 2008;73(1):158–170.

13. Shigesada N, Kawasaki K, Teramoto E. Spatial segregation of interacting species. Journal of Theoretical Biology. 1979;79(1):83–99.

14. Shigesada N, Kawasaki K, Teramoto E. Traveling periodic waves in heterogeneous environments. Theoretical Population Biology. 1986;30(1):143–160.

15. Shigesada N, Kawasaki K. Biological invasions: theory and practice. Oxford University Press, UK; 1997.

16. Hastings A, Cuddington K, Davies KF, Dugaw CJ, Elmendorf S, Freestone A, et al. The spatial spread of invasions: new developments in theory and evidence. Ecology Letters. 2005;8(1):91–101.

17. Schreiber SJ, Lloyd-Smith JO. Invasion dynamics in spatially heterogeneous environments. The American Naturalist. 2009;174(4):490–505.

18. Dewhirst S, Lutscher F. Dispersal in heterogeneous habitats: thresholds, spatial scales, and approximate rates of spread. Ecology. 2009;90(5):1338–1345.

19. Wolf M, Weissing FJ. Animal personalities: consequences for ecology and evolution. Trends in ecology & evolution. 2012;27(8):452–461.

20. Sih A, Cote J, Evans M, Fogarty S, Pruitt J. Ecological implications of behavioural syndromes. Ecology letters. 2012;15(3):278–289.

21. Chapple DG, Simmonds SM, Wong BB. Can behavioral and personality traits influence the success of unintentional species introductions? Trends in Ecology & Evolution. 2012;27(1):57–64.

22. Carere C, Gherardi F. Animal personalities matter for biological invasions. Trends in ecology & evolution. 2013;28(1):5–6.

23. Frankel NW, Pontius W, Dufour YS, Long J, Hernandez-Nunez L, Emonet T. Adaptability of non-genetic diversity in bacterial chemotaxis. Elife. 2014;3:e03526.

24. Dufour YS, Fu X, Hernandez-Nunez L, Emonet T. Limits of feedback control in bacterial chemotaxis. PLoS computational biology. 2014;10(6):e1003694.

25. Dufour YS, Gillet S, Frankel NW, Weibel DB, Emonet T. Direct correlation between motile behavior and protein abundance in single cells. PLoS computational biology. 2016;12(9):e1005041.

26. Fogarty S, Cote J, Sih A. Social personality polymorphism and the spread of invasive species: a model. The American Naturalist. 2011;177(3):273–287.

27. Ben-Jacob E, Cohen I, Levine H. Cooperative self-organization of microorganisms. Advances in Physics. 2000;49(4):395–554.

28. Keller EF, Segel LA. Traveling bands of chemotactic bacteria: a theoretical analysis. Journal of theoretical biology. 1971;30(2):235–248.

29. Lin TC, Wang ZA. Development of traveling waves in an interacting two-species chemotaxis model. Discrete Continuous Dynamical Systems Series A. 2014;34(7):2907–2927.

30. Emako C, Gayrard C, Buguin A, de Almeida LN, Vauchelet N. Traveling pulses for a two-species chemotaxis model. PLoS computational biology. 2016;12(4):e1004843.

31. Veening JW, Smits WK, Kuipers OP. Bistability, epigenetics, and bet-hedging in bacteria. Annu Rev Microbiol. 2008;62:193–210.

32. Kussell E, Leibler S. Phenotypic diversity, population growth, and information in fluctuating environments. Science. 2005;309(5743):2075–2078.

33. Wolf DM, Vazirani VV, Arkin AP. Diversity in times of adversity: probabilistic strategies in microbial survival games. Journal of theoretical biology. 2005;234(2):227–253.

34. Wolf DM, Vazirani VV, Arkin AP. A microbial modified prisoner’s dilemma game: how frequency-dependent selection can lead to random phase variation. Journal of theoretical biology. 2005;234(2):255–262.

35. Solopova A, van Gestel J, Weissing FJ, Bachmann H, Teusink B, Kok J, et al. Bet-hedging during bacterial diauxic shift. Proceedings of the National Academy of Sciences. 2014;111(20):7427–7432.

36. Stumpf MP, Laidlaw Z, Jansen VA. Herpes viruses hedge their bets. Proceedings of the National Academy of Sciences. 2002;99(23):15234–15237.

37. Rouzine IM, Weinberger AD, Weinberger LS. An evolutionary role for HIV latency in enhancing viral transmission. Cell. 2015;160(5):1002–1012.

38. Childs DZ, Metcalf C, Rees M. Evolutionary bet-hedging in the real world: empirical evidence and challenges revealed by plants. Proceedings of the Royal Society of London B: Biological Sciences. 2010; p. rspb20100707.

39. Hidalgo J, de Casas RR, Muñoz MÁ. Environmental unpredictability and inbreeding depression select for mixed dispersal syndromes. BMC evolutionary biology. 2016;16(1):71.

40. Hopper KR. Risk-spreading and bet-hedging in insect population biology. Annual review of entomology. 1999;44(1):535–560.

41. Kelly Jr JL. A new interpretation of information rate. In: The Kelly Capital Growth Investment Criterion: Theory and Practice. World Scientific; 2011. p. 25–34.

42. Fernholz R, Shay B. Stochastic portfolio theory and stock market equilibrium. The Journal of Finance. 1982;37(2):615–624.

43. Smith JM. Evolution and the Theory of Games. In: Did Darwin Get It Right? Springer; 1988. p. 202–215.

44. Nowak MA. Evolutionary dynamics. Harvard University Press; 2006.

45. Harmer GP, Abbott D. Game theory: Losing strategies can win by Parrondo’s paradox. Nature. 1999;402(6764):864.

46. Parrondo JM, Harmer GP, Abbott D. New paradoxical games based on Brownian ratchets. Physical Review Letters. 2000;85(24):5226.

47. de Jong IG, Haccou P, Kuipers OP. Bet hedging or not? A guide to proper classification of microbial survival strategies. Bioessays. 2011;33(3):215–223.

48. Williams PD, Hastings A. Paradoxical persistence through mixed-system dynamics: towards a unified perspective of reversal behaviours in evolutionary ecology. Proceedings of the Royal Society of London B: Biological Sciences. 2011; p. rspb20102074.

49. Comins HN, Hamilton WD, May RM. Evolutionarily stable dispersal strategies. Journal of theoretical Biology. 1980;82(2):205–230.

50. Hamilton WD, May RM. Dispersal in stable habitats. Nature. 1977;269(5629):578.

51. Jansen VA, Yoshimura J. Populations can persist in an environment consisting of sink habitats only. Proceedings of the National Academy of Sciences. 1998;95(7):3696–3698.

52. Hidalgo J, Pigolotti S, Munoz MA. Stochasticity enhances the gaining of bet-hedging strategies in contact-process-like dynamics. Physical Review E. 2015;91(3):032114.

53. Rajon E, Venner S, Menu F. Spatially heterogeneous stochasticity and the adaptive diversification of dormancy. Journal of evolutionary biology. 2009;22(10):2094–2103.

54. Den Boer PJ. Spreading of risk and stabilization of animal numbers. Acta biotheoretica. 1968;18(1–4):165–194.

55. Hufton PG, Lin YT, Galla T. Phenotypic switching of populations of cells in a stochastic environment. Journal of Statistical Mechanics: Theory and Experiment. 2018;2018(2):023501.

56. Ashcroft P, Altrock PM, Galla T. Fixation in finite populations evolving in fluctuating environments. Journal of The Royal Society Interface. 2014;11(100):20140663.

57. Hufton PG, Lin YT, Galla T, McKane AJ. Intrinsic noise in systems with switching environments. Physical Review E. 2016;93(5):052119.

58. Van Saarloos W. Front propagation into unstable states. Physics reports. 2003;386(2–6):29–222.

59. Cencini M, Lopez C, Vergni D. Reaction-diffusion systems: front propagation and spatial structures. In: TheKolmogorov Legacy in Physics. Springer; 2003. p. 187–210.

60. Korolev KS, Avlund M, Hallatschek O, Nelson DR. Genetic demixing and evolution in linear stepping stone models. Reviews of modern physics. 2010;82(2):1691.

61. Dornic I, Chaté H, Munoz MA. Integration of Langevin equations with multiplicative noise and the viability of field theories for absorbing phase transitions. Physical review letters. 2005;94(10):100601.

62. Moro E. Numerical schemes for continuum models of reaction-diffusion systems subject to internal noise. Physical Review E. 2004;70(4):045102.

63. Brunet E, Derrida B. Effect of microscopic noise on front propagation. Journal of Statistical Physics. 2001;103(1–2):269–282.

64. Moro E. Internal fluctuations effects on Fisher waves. Physical Review Letters. 2001;87(23):238303.

65. Hofbauer J, Sigmund K. Evolutionary games and population dynamics. Cambridge university press; 1998.

66. Kapur J, Rani A. Testing the consistency of given values of a set of moments of a probability distribution. J Bihar Math Soc. 1995;16:51–63.

67. Jones SE, Lennon JT. Dormancy contributes to the maintenance of microbial diversity. Proceedings of the National Academy of Sciences. 2010;107(13):5881–5886.

68. Møller AP, Garamszegi LZ. Between individual variation in risk-taking behavior and its life history consequences. Behavioral Ecology. 2012;23(4):843–853.

69. Xue B, Leibler S. Evolutionary learning of adaptation to varying environments through a transgenerational feedback. Proceedings of the National Academy of Sciences. 2016;113(40):11266–11271.

70. Pigolotti S, Benzi R. Selective advantage of diffusing faster. Physical review letters. 2014;112(18):188102.

71. Pigolotti S, Benzi R. Competition between fast-and slow-diffusing species in non-homogeneous environments. Journal of theoretical biology. 2016;395:204–210.

72. Gueudré T, Dobrinevski A, Bouchaud JP. Explore or exploit? A generic model and an exactly solvable case. Physical review letters. 2014;112(5):050602.

73. Gandhi SR, Yurtsev EA, Korolev KS, Gore J. Range expansions transition from pulled to pushed waves as growth becomes more cooperative in an experimental microbial population. Proceedings of the National Academy of Sciences. 2016;113(25):6922–6927.

74. Birzu G, Hallatschek O, Korolev KS. Fluctuations uncover a distinct class of traveling waves. Proceedings of the National Academy of Sciences. 2018;115(16):E3645-E3654.

75. Baym M, Lieberman TD, Kelsic ED, Chait R, Gross R, Yelin I, et al. Spatiotemporal microbial evolution on antibiotic landscapes. Science. 2016;353(6304):1147–1151.

76. Müller MJ, Neugeboren BI, Nelson DR, Murray AW. Genetic drift opposes mutualism during spatial population expansion. Proceedings of the National Academy of Sciences. 2014;111(3):1037–1042.

